# A Universal Nonparametric Event Detection Framework for Neuropixels Data

**DOI:** 10.1101/650671

**Authors:** Hao Chen, Shizhe Chen, Xinyi Deng

## Abstract

Neuropixels probes present exciting new opportunities for neuroscience, but such large-scale high-density recordings also introduce unprecedented challenges in data analysis. Neuropixels data usually consist of hundreds or thousands of long stretches of sequential spiking activities that evolve non-stationarily over time and are often governed by complex, unknown dynamics. Extracting meaningful information from the Neuropixels recordings is a non-trial task. Here we introduce a general-purpose, graph-based statistical framework that, without imposing any parametric assumptions, detects points in time at which population spiking activity exhibits simultaneous changes as well as changes that only occur in a subset of the neural population, referred to as “change-points”. The sequence of change-point events can be interpreted as a footprint of neural population activities, which allows us to relate behavior to simultaneously recorded high-dimensional neural activities across multiple brain regions. We demonstrate the effectiveness of our method with an analysis of Neuropixels recordings during spontaneous behavior of an awake mouse in darkness. We observe that change-point dynamics in some brain regions display biologically interesting patterns that hint at functional pathways, as well as temporally-precise coordination with behavioral dynamics. We hypothesize that neural activities underlying spontaneous behavior, though distributed brainwide, show evidences for network modularity. Moreover, we envision the proposed framework to be a useful off-the-shelf analysis tool to the neuroscience community as new electrophysiological recording techniques continue to drive an explosive proliferation in the number and size of data sets.

## 1. Introduction

Electrophysiological recording techniques have become more sophisticated, incorporating simultaneous spiking data from more neurons across multiple brain regions. In particular, hair-thin probes densely packed with hundreds of recording sites, called Neuropixels, can record spiking activity from hundreds or even thousands of cells [1, 2]. The combined high temporal resolution and broad spatial coverage of these probes offer a new picture of the coordinated activity in the brain. For example, the International Brain Laboratory, a collaboration of 21 laboratories, has recently pledged to include Neuropixels as one of three complementary recording techniques to reveal how and where decision-related signals change over time and interact across regions as an animal commits to a choice [3].

Such large-scale, high-density recordings of neural activity pose new analysis challenges that must be resolved for the effective use of these data in order to answer questions about how the whole brain works together [4, 5, 6, 7, 8]. Neuropixels probes record simultaneously and continuously from thousands of neurons for long stretches of time. Even though initial processing steps on these data can divide the spike data into ~ 10 brain regions [2], each brain region in general still contains hundreds of sequences of spiking activities and these sequences are correlated in a complex way (see Figure 1). In addition, these high-dimensional spiking sequences evolve non-stationarily over time, making the understanding of these sequences even more difficult. Currently, there is no working model that can handle these sequences in all brain regions effectively. Here we present a new statistical framework that addresses these challenges to help understand and draw scientific conclusions from large-scale neural population recordings with Neuropixels probes.

**Fig. 1.**
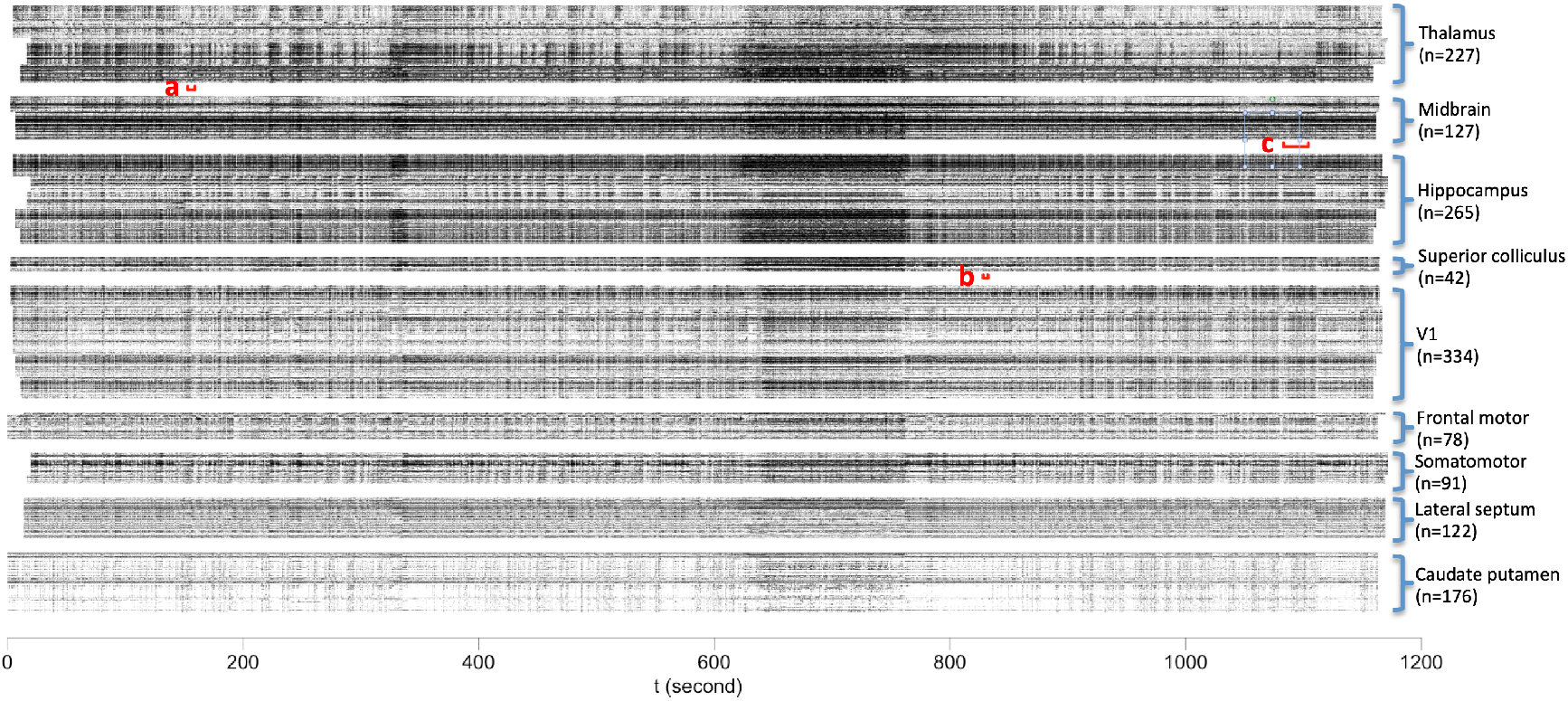
Eight-probe Neuropixels recordings across nine brain regions in a mouse awake in darkness during spontaneous behavior. Each row corresponds to one neuron. The neurons are grouped by brain regions with the number of neurons listed in the parentheses. Three intervals in different brain regions, a, b and c, are used to illustrate our method with details in Figure 2. Brain region labels were provided by Steinmetz et al. at doi.org/10.25378/janelia.7739750.

### 1.1. Our Contributions

In this work, we provide a practical procedure to process and analyze the Neuropixels data efficiently. Specifically, we propose a nonparametric framework that identifies the change-points of a multivariate point process, where a change-point is defined as the time point when the probabilistic law of the point process abruptly changes (see Figure 2). To put it in the context of Neuropixels data, an immediate product of the proposed procedure is a series of time points when the neural activity significantly changes in the long sequence of spiking activities for each brain region. These time points divide the long sequence into stationary segments to facilitate more detailed follow-up investigations. Moreover, the series of change-points for each brain region serves as a fingerprint of the region, through which we can infer the relationship among the regions, including clustering regions into different groups.

**Fig. 2.**
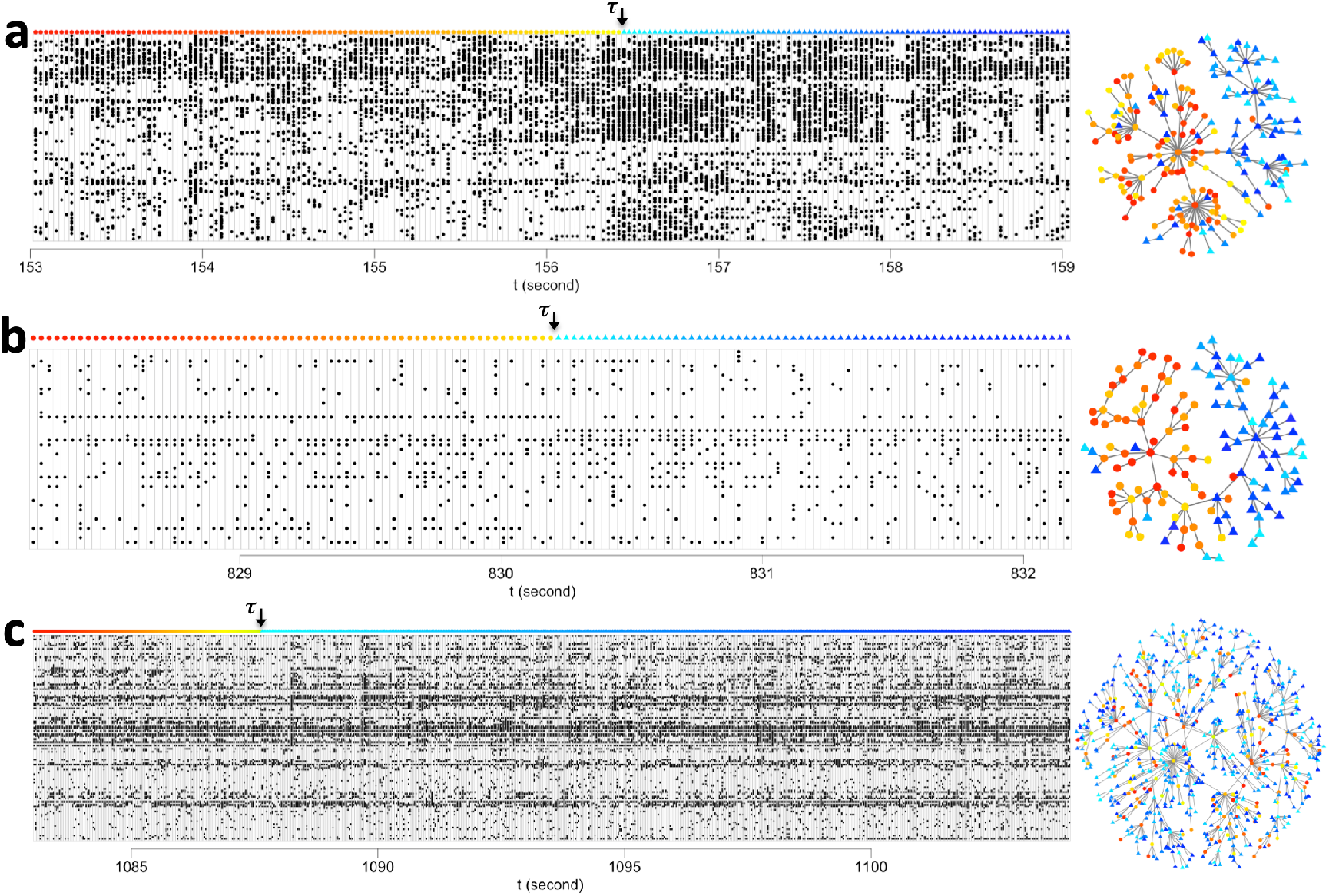
An illustration of the change-point detection method applied to Neuropixels data. The time interval and brain region corresponding to each panel, a, b and c, in the whole data sequence are denoted in red bracket in Figure 1. Here, the spike train data are plotted with the estimated change-point denoted by *τ*. For each interval, the similarity graph constructed in the change-point detection method is shown on the right with each dot corresponding to the vector representing neuron-spiking information of all neurons in that brain region at one time. We see that the method could detect the overall intensity changes (interval a), it could also detect the change when the overall intensity is similar while the active neurons change (interval b), or the covariance matrix of the spiking vector changes (interval c). Detailed description of the procedure is provided in Section 2, and summary statistics for the three intervals are provided in Table 1.

Our main contribution is providing a new angle for initial exploratory analysis of the Neuropixels data. As a compliment to existing exploratory summary statistics such as the overall firing intensity, peri-stimulus time histogram, correlogram, spike-triggered average, and low-dimensional Gaussian-process factors [9], we provide the change-points as a low-dimensional summary of the activities of a large neural population. Because of the simplicity of the proposed pipeline, change-points can be easily detected using a computationally-efficient algorithm and require few assumptions. All these are made possible because of the technical advantages of our proposal as follows.

i. **A statistically-principled nonparametric approach:** Our proposed pipeline builds upon a graph-based change-point detection framework [10, 11], which is rooted in the statistical theory of detecting lack of homogeneity in a sequence of observations. Moreover, our method imposes no distributional or parametric assumptions on the data. The absence of such assumptions makes the proposed method especially appealing for exploratory analysis of complex neural activity data, where one may want to avoid artifactual discovery due to implicit assumptions in the methods.
ii. **Detecting various change types in high dimensions:** Event detection is challenging as the dimensionality increases, commonly known as the *curse of dimensionality*. The proposed pipeline accounts for effects resulting from the curse of dimensionality to effectively detect various types of change-points for data in high dimensions (see Figure 2 for an illustration), which makes it suitable for handling activities of large neural ensembles in Neuropixels recordings.
iii. **An easy-to-use off-the-shelf tool:** We provide easy-to-compute summary statistics – the change-points – of neural activities in each brain region over a long recording. As demonstrated in Figure 3, the detected change-points reveals the footprint of activities in each brain region. Furthermore, the proposed pipeline incurs low computational cost and has been implemented in a software package. As a result, the proposed method can be incorporated, at little cost, in the routine of initial assessment of Neuropixels data.

**Fig. 3.**
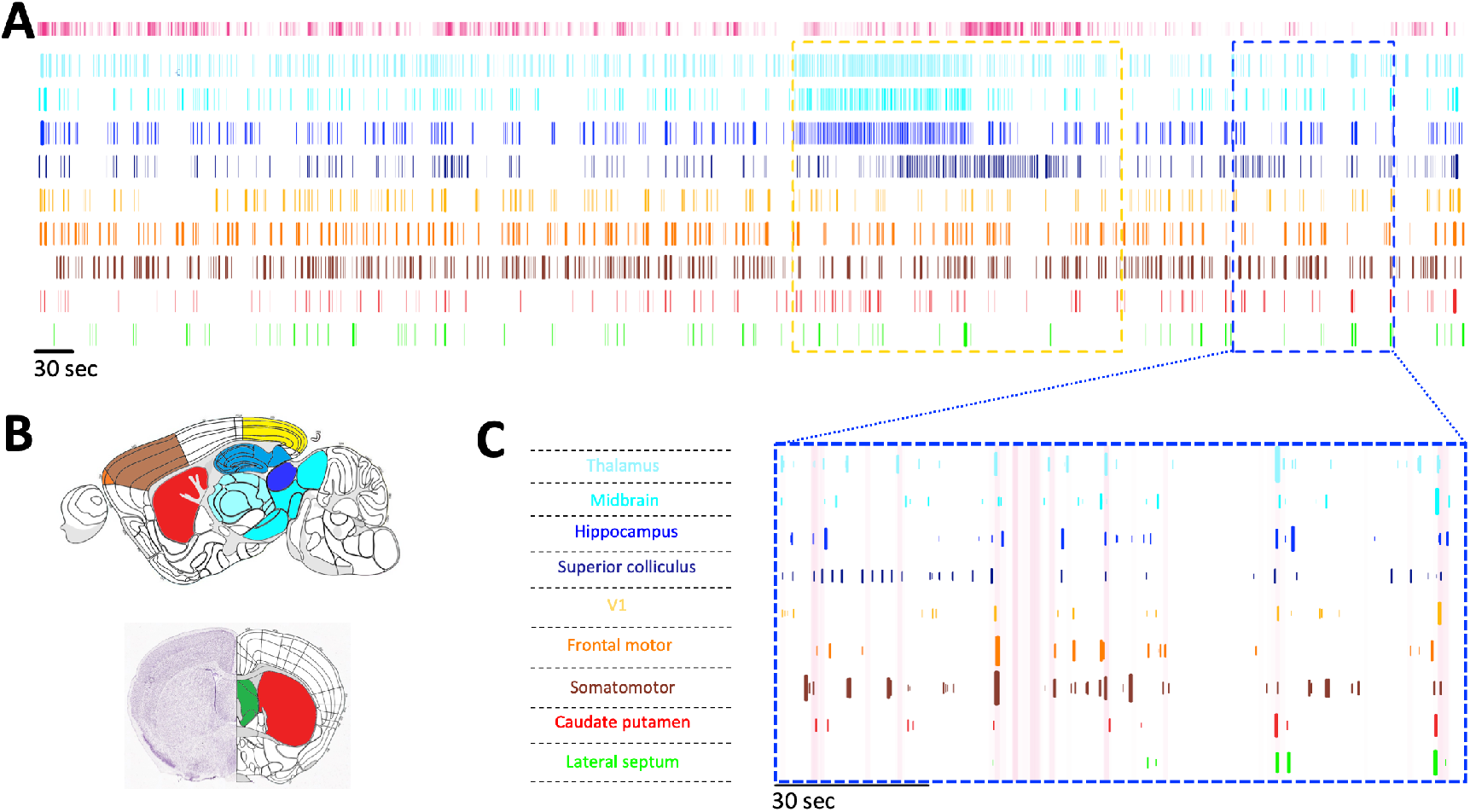
Change-points detected in the Neuropixels recordings using the proposed procedure. **A**, Detected change-points for individual brain regions. Each vertical colored line is a change-point with the width of the line reflects the significance of the change-point; each row plots the change-points for an individual brain region. *Top insert*: Face motion energy is plotted in pink, calculated as the variances of the top 50 singular values provided by [2]. Change-point dynamics display similarities among some brain regions and can be visually clustered in to four modules (colored-coded by blue hues, yellow hues, red, and green, respectively). Blue hues: thalamus, hippocampal formation, midbrain, superior colliculus. Yellow hues: frontal motor cortex, somato-motor cortex, V1. Red: caudate putamen. Green: lateral septal nucleus. Change-point dynamics across modules differ from “internal-pathway driven” (yellow dashed outline) and “behaviorally aligned” phases (blue dashed outline). **B**, Brain atlases colored with the coloring scheme used in panel A. Reference atlases for the sagittal and coronal views of the mouse brain obtained from http://atlas.brain-map.org/. **C**, Zoom-in to an example period (blue dashed outline) where the change-points are aligned across brain regions to face motion energy (pink background). Here, the height and width of a change-point reflect its significance.

### 1.2. Outline

In the following, we discuss in details the proposed method designed for dealing with long high-dimensional sequences with multiple change-points (Section 2). We apply the method to a Neuropixels recording collected during spontaneous behavior of an awake mouse in complete darkness, and the results are presented and discussed in Section 3. We conclude the paper with discussion in Section 4.

## 2. Multiple Event Detection in a Long High-Dimensional Sequence

In this section, we first briefly review the graph-based change-point detection framework developed in [10, 11] in Section 2.1. The existing framework focuses on sequences with at most one change-point or one changed-interval, which is ill-suited for analyzing Neuropixels data. In Section 2.2, we present an effective way to detect multiple change-points in a high-dimensional sequence. The concrete algorithm that can be directly applied to the Neuropixels data is stated in Section 2.3.

### 2.1. Graph-Based Change-Point Detection

Let **y**_1_, **y**_2_,…, **y**_*T*_ be the data sequence, where **y**_*t*_ is an *n*-dimensional vector that we refer to as one *observation* in what follows. There possibly exists a time *τ* such that **y**_*t*_ has one distribution for *t* ≤ *τ* and another distribution for *t* > *τ*. In these works, the authors adapted graph-based two-sample tests [12, 13, 14] to the scan statistic framework: Each *t* divides the observations into two groups, {**y**_1_,…, **y**_*t*_} and {**y**_*t*+1_,…, **y**_*T*_}, and a graph-based two-sample test is conducted to test whether these two groups of observations are from the same distribution or not. Then, the maximum of the scan statistics over *t* is used as the test statistic.

The graph-based two-sample tests are tests that are based on a similarity graph constructed on all observations. The similarity graph can be any given graph that reflects the similarity between observations [15]. More generally, it can be constructed based on a similarity measure through a certain criterion, such as a minimum spanning tree (MST) [12], which is a tree connecting all observations with the total distance across edges minimized, or a nearest neighbor graph where each observation connects to its nearest neighbor [16, 17]. Let *G* be the similarity graph on all observations in the sequence. The graph-based two-sample tests are based on two basic quantities computed from the graph. Let *g_i_*(*t*) = *I*_*i*>*t*_, where *I_A_* is an indicator function that takes value 1 if event *A* is true and 0 otherwise. Then the two quantities are:

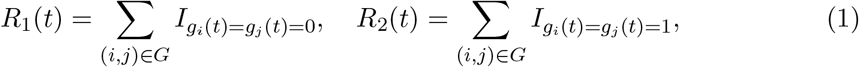

where *R*_1_(*t*) counts the number of edges connecting observations before time *t*, and *R*_2_(*t*) counts the number of edges connecting observations after *t*. A number of scan statistics were explored in [10, 11]. Here, we focus on the generalized edge-count scan statistic as it takes into account the *curse of dimensionality* and works properly and effectively for high-dimensional data in detecting various types of events [13]:

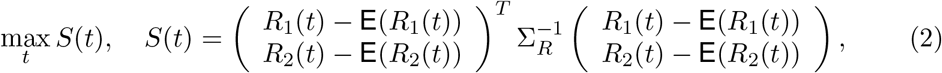

where Σ_*R*_ = **Var**((*R*_1_(*t*), *R*_2_(*t*))^*T*^). Since there is no distributional assumption for the data, the expectation and variance are defined on the permutation null distribution, which places probability 1/*T*! on each of the *T*! permutations of {**y**_1_, **y**_2_,…, **y**_*T*_}. Exact analytic formulas for **E**(*R*_1_(*t*)), **E**(*R*_2_(*t*)) and **Var**((*R*_1_(*t*), *R*_2_(*t*))^*T*^) are provided in [11] to ease the computation of the scan statistics. In addition, the authors provided analytic formulas to approximate the permutation *p*-value of the scan statistic, for 1 < *T*_0_ < *T*_1_ < *T*,

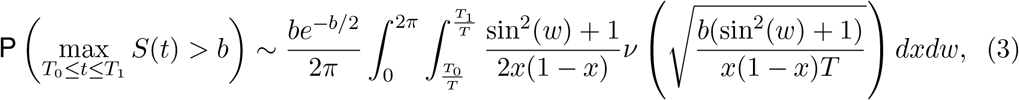

where 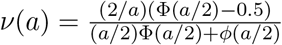 with Φ(·) and *ϕ*(·) the standard normal cumulative density function and probability density function, respectively. Based on this analytic expression, one can easily determine the threshold for the scan statistic at any type I error rate under the permutation null, allowing fast detection. Furthermore, it is possible to determine the types of the change-points based on the values of *R*_1_(*t*) and *R*_2_(*t*). As a concrete example, in Table 1, we provide the test statistics and the corresponding *p*-values for the three change-points shown in Figure 2.

**Table 1.** Summary statistics of the change-points in Figure 2. Here *S*(*t*) is provided in (2) and *p*-value is computed through (3), 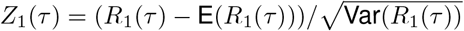 and 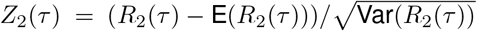 are the standardized edge counts (*Z*_1_(*τ*) and *Z*_2_(*τ*) roughly follow the standard normal distribution, respectively). In intervals a and b, the differences between the vector of the intensity of the neuron’s spiking rate before and after *τ* dominates the change, causing the observations before *τ* and after *τ* well separated in the similarity graph. As a result, the values of *Z*_1_(*τ*) and *Z*_2_(*τ*) are both large and *S*(*τ*) large. In interval a, the overall intensity after *τ* increases; while in interval b, the overall intensity after *τ* is similar to that before *τ* but the set of neurons that are active differ. In interval c, the covariance matrix of the neuron spiking intensity before and after *τ* dominates the change and the determinant of the covariance matrix after *τ* is larger causing the observations before *τ* are mainly hubs and the observations after *τ* are mainly leafs in the similarity graph. As a result, the value of *Z*_1_(*τ*) is large, the value of *Z*_2_(*τ*) is negatively large, and *S*(*τ*) large.

| Region | $\max_t S(t)$ | $p$-value | $Z_1(\tau)$ | $Z_2(\tau)$ |
| --- | --- | --- | --- | --- |
| a | 125.8 | $6.1 \times 10^{-26}$ | 3.0 | 3.1 |
| b | 92.1 | $8.7 \times 10^{-19}$ | 4.8 | 4.7 |
| c | 90.3 | $4.7 \times 10^{-18}$ | 9.2 | -5.5 |

#### Remark 1.

*One may notice that the permutation null is stronger than the null hypothesis that the spike trains are stationary. In particular, stationarity of point processes allows the existence of temporal dependence, whereas the permutation null prohibits any form of dependence in the sequence. In fact, an extension of the presented procedure that allows for short-term temporal dependence has been developed [18], where the circular block permutation replaces the plain-vanilla permutation for constructing the null distribution of test statistics. However, the detection framework using circular block permutation demands much higher computational cost than the presented detection method using the plain-vanilla permutation. Therefore, in this work, we choose to present the change-point detection framework using the permutation null that allows for quick exploration of big data such as Neuropixels recordings. We note that, if one desires precise control of the type I error rate, it is recommended to use the circular block permutation method to refine the detected change-points*.

### 2.2. Multiple Change-Point Detection

In Neuropixels data, there are in general multiple change-points and the number of change-points could even be proportional to the duration of recordings. The existing method is thus infeasible for analyzing long sequences of Neuropixels recordings. In the following, we present a set of practical steps to detect multiple change-points in the sequence effectively.

#### 2.2.1. Finding an Initial Set of Candidate Change-Points

We adapt the binary segmentation approach [19] to find the initial set of candidate change-points when there are multiple change-points. The procedure is as follows: We apply the graph-based change-point detection approach to the whole sequence to find the first change-point. This change-point divides the whole sequence into two parts and we apply the approach to each sub-sequence. This process keeps going until no more change-point can be found. Note that this step only gives us an initial set of candidate change-points and these change-points are refined in the later iterative steps.

There are a few other popular methods for finding multiple change-points, such as circular binary segmentation [20, 21] and wild binary segmentation [22]. All these approaches usually give sub-optimal estimates of the change-points when there are multiple change-points. However, to get the optimal estimates of change-points, one needs to consider all possible choices of the locations of the change-points, that is 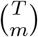 where *m* is the true number of change-points, which would be computationally prohibitive when *m* is not too small. Here, we choose the practical binary segmentation approach that is easy and fast to implement, which is essential in analyzing long Neuropixels recordings.

When the length of the sequence *T* is large, we chop the whole sequence into ⌈*T/c*_1_⌉ sub-sequences, each with length *c*_1_ + *c*_2_ and consecutive sub-sequences overlapped by the length of *c*_2_. We then apply the binary segmentation approach to each of these sub-sequences. Here, *c*_1_ and *c*_2_ are two constants. The procedure is not sensitive to the choice of *c*_1_ and *c*_2_, and a reasonable choice can be *c*_1_ = 1,000 and *c*_2_ = 200. Using the sub-sequence detection may potentially include more false discoveries, which will be adjusted in the later refinement steps, but it significantly reduces the time complexity of the initial detection. Let *n* be the dimension of the sequence. The time complexity of applying the binary segment approach to the entire sequence using the graph-based statistics is typically *O*(*nT*^2^ log *T*) and under the worst-case scenario *O*(*nT*^3^); while with this extra chopping step, the time complexity becomes *O*(*n*(*c*_1_ + *c*_2_)^3^*T/c*_1_) = *O*(*nT*) under the worst-case scenario. Hence, without further specification, this chopping step is employed when *T* is large.

#### 2.2.2. Change-Point Refinement

Suppose, after *i*^th^ iteration, *i* = 0,1, 2,…, there are *K*^(3*i*)^ candidate change-points and we denote them by 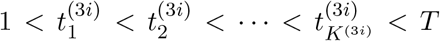. For notation simplicity, we let 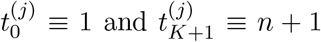, ∀*j*. Due to the limitation in the earlier steps, it could be that the estimated change-point is off from the true change-point. In this step, we first refine the estimates of the change-points. In particular, for each change-point 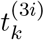, *k* ∈ {1,…, *K*^(3*i*)^}, the graph-based change-point detection approach is applied to the interval 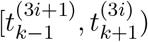 to refine the estimate, and the refined change-point is denoted by 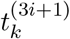.

Given that the estimates of the change-points are altered, we further check if there is any change-point in each sub-interval 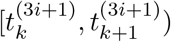, *k* ∈ {0,…, *K*^(3*i*)^}, with the threshold *b* determined by (3) with “*T*” in the equation replaced by the number of observations in this interval 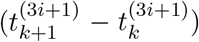 and the probability being controlled at *α/K*^(3*i*)^, where *α* is a user-defined error rate, such as 0.01. Let *K*^(3*i*+2)^ be the number of candidate change-points after the re-searching and we denote these change-points by 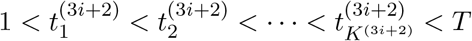.

After the previous steps, some of the candidate change-points might correspond to the same event. We thus prune the change-points, that is, 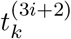, *k* ∈ {1,…, *K*^(3*i*+2)^}, is kept only if the observations in the time interval 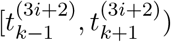 is significantly non-homogeneous. In particular, we use the Benjamini-Yekutieli procedure [23] to control the false discover rate at *α*.

### 2.3. Algorithm

#### Algorithm 1

~~~
1: **Initialization:** use the binary segmentation approach combined with the scan statistic on *S* to find the initial set of change-points.
2: **while** not convergence or reach the maximum number of iterations **do**
3:   Refine the estimates to the time points of the current set of change-points;
4:   Find possible extra change-points in each sub-interval;
5:   Prune out insignificant change-points with false discovery rate set at *α*;
6: **end**
7: Compute the significance level for the final set of change-points.
~~~

Algorithm 1 summarizes the steps described in Section 2.2. The convergence rule is that there is no change in the estimated change-points. It may take some time to converge, and one can set the maximum number of iterations. In practice, twenty iterations yield reasonably good estimates of the change-points. The computing complexity of the proposed algorithm is provided in Theorem 1. The proof is deferred to the Appendix.

#### Theorem 1.

*For a sequence of length T and dimension n, the time complexity of Algorithm 1 is O*(*nT*) *when* 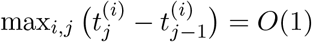, *and is O*(*nT*^2^) *under the worst scenario when* 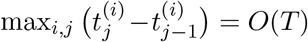. *The space complexity of Algorithm 1 is O*(*nT*) *when* 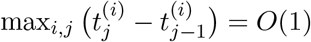, *and is O*(*T* max(*n, T*)) *under the worst scenario*.

## 3. Application to Neuropixels Recordings

In this section, we present an analysis of an eight-probe Neuropixels data set recorded during spontaneous behavior of a mouse awake in darkness [24] using the proposed change-point detection procedure. The recordings have been spike-sorted and preprocessed by its providers [2]. The data set contains spike trains of 1,462 neurons simultaneously recorded for a period of around 20 minutes. These 1,462 neurons have been grouped into 9 brain regions and the full recording has been discretized into 39,053 intervals of 30ms. This data set can be downloaded from https://janelia.figshare.com/articles/Eight-probe_Neuropixels_recordings_during_spontaneous_behaviors/7739750. The entire data set is plotted in Figure 1 in Section 1.1. A behavioral data set, also provided by [2], was recorded concurrently with neural activity and contained the processed version of a mouse face movie.

We show in panel A of Figure 3 the detected change-points over the full recording period with false discovery rate *α* in Algorithm 1 set to be 0.01. The running time for these 9 brain regions ranges from 10 minutes to 1 hour on a personal laptop (the Macbook 2015, 1.2 GHz Intel Core M) depending on the number of neurons in the sequence and how sparse the change-points are. The algorithm could be much expedited through parallelization for most of its steps, not done in the current implementation.

In the following, we focus on two example time windows, within the full recording period, where the change-point patterns show biologically interesting dynamics.

### Case study 1: Internal pathways hinted by the change-point patterns

Outlined with yellow dashed lines in panel A of Figure 3, we observe a period where the patterns of the detected change-points exhibit striking similarity among brain regions, which suggests possible coordination and/or regulation across brain regions. Recall that a change-point represents an abrupt change of neural activity. Therefore, the pattern of change-points reflects the ongoing process in the corresponding brain region. For instance, a period of dense change-points indicates that the corresponding brain region is in a highly-dynamic state, whereas the absence of change-points represents that the neural activity is rather stationary. One notable feature of the change-points within the yellow dashed lines is that there is clearly a cluster of high density change-points in thalamus, midbrain, hippocampus, and superior colliculus. The clusters of change-points onset around the same time in thalamus, midbrain and hippocampus, indicating that this is either a coordinated response or a result of common input. The clear time delay in the superior colliculus may indicate that superior colliculus is downstream in the information flow for this particular event. Furthermore, these clusters of change-points are followed by a period of high face motion energy. It is likely that the face motion is a consequence of this coordinated activity. Of note, these four brain regions showing similar patterns in change-points corroborate known connectivity in the mouse nervous system (see, among others, [25, 26, 27, 28]).

### Case study 2: Behaviorally aligned change-points

Outlined with blue dashed lines in panel A of Figure 3 with panel C a zoom-in display, we observe a period where the detected change-points align well with onsets of high face motion energy. Note that the change-points are temporally-local features; that is, the timing of a change-point depends locally on the neural activity before and after the change-point. As a result, the change-point detection has high temporal resolution, which reveal possible connections between the neural activity and the behavior of the animal as shown in panel C.

It is important to point out that the results above should be interpreted as *initial findings* from the proposed change-point detection method. Further investigations are warranted to understand the underlying mechanisms of the detected change points and their relationships with behavior.

## 4. Discussion

We have introduced a general-purpose, graph-based framework for change-point detection suitable for long sequences of large-scale neural recordings. We demonstrate the effectiveness of our method in the analysis of a Neuropixels recording across multiple brain regions during spontaneous behavior of an awake mouse in complete darkness. Our analyses show that neural activity underlying spontaneous behavior, though distributed brainwide [2], suggests evidence for network modularity. Our observations corroborate previous studies on anatomical and functional networks of the mouse brain [25, 26, 27, 28] and could lead to spatially constrained predictions about brain function that may be tested in terms of lesions, evoked responses, and dynamic patterns of neural activities.

An important observation is that the graph-based approach, not only easy and flexible to implement with low computational complexity, has desirable power when compared with existing parametric tests in moderate and high dimensions [15, 10, 11]. The proposed procedure provides an off-the-shelf analytical tool that is simple to integrate into existing neural data analysis pipelines. We envision the proposed framework to be increasingly useful to the neuroscience community as new electrophysiological recording techniques continue to drive an explosive proliferation in the number and size of data sets involving large numbers of neurons across multiple brain regions.

## Acknowledgments

The authors thank Kenneth Kay and Helen Xun Hou for helpful discussions. HC was supported by NSF DMS-1513653. XD additionally thanks Liam Paninski for organizational support during her time on this project. XD was supported by the NSF NeuroNex Award (DBI-1707398), the Gatsby Charitable Foundation, and the Simons Foundation (SCGB 543023).

## A. Proof of Theorem 1

For the graph-based change-point detection approach applying to a sequence of length *T*_0_ and dimension *n*, its time complexity is bottlenecked by the computation of pairwise distances among the observations, i.e., 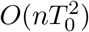. The binary segmentation approach in the initialization step would typically add a factor of log(*T*_0_) in time complexity, but a factor of *O*(*T*_0_) under the worst-case scenario. After adopting the extra chopping step elaborated in Section 2.2.1, the time complexity for the initialization step becomes *O*(*nT*). In the refinement step, the time complexity for applying the graph-based change-point detection approach to time interval [*t_j_,t*_*j*+1_) is *O*(*n*(*t*_*j*+1_ − *t_j_*)^2^). When 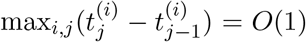, the time complexity of the refinement step is *O*(*nT*). Note that 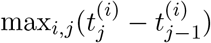 can be as large as *T*, leading to the worst-case complexity of *O*(*nT*^2^). For the space complexity, it is bottlenecked by the size of the data *O*(*nT*) or the pairwise distance matrices during the data analysis step, which could be as large as *O*(*T*^2^) when 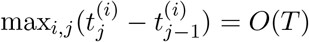.

